# Gut Microbial Metabolic Disorder in Depression: Insights from Computational Modeling and Mediation Analysis

**DOI:** 10.1101/2024.10.10.617513

**Authors:** Yuchen Zhang, Wenkai Lai, Meiling Wang, Shirong Lai, Qing Liu, Qi Luo, Fenglong Yang

## Abstract

**Background:** Depression is a common mental disorder worldwide, and its pathogenesis remains incompletely understood. However, increasing evidence suggests that gut microbiota play a significant role in the development of depression through the gut-brain-microbiota axis. However, due to the substantial individual variability in gut microbiota, while metabolic functions and metabolites between individuals are relatively similar, analyzing functional profiles and metabolic products yields more accurate results.

**Results:** In this study, gut microbiota abundance data were collected from 354 depression patients and 5575 healthy individuals in the GMrepo v2 database. After matching for age and BMI, 271 paired samples from each group were retained. Alpha diversity analysis revealed significant differences between the two groups in the Observed, Shannon, and Chao1 indices, while beta diversity analysis indicated only subtle differences in gut microbiota composition. LEfSe analysis identified 23 differential species, with 18 enriched in the healthy group and 5 in the depression group. Further alpha diversity analysis of reaction abundance showed significant differences in the Observed and Chao1 indices, while beta diversity analysis did not reveal significant differences in reaction abundance. Differential and enrichment analyses identified 89 reactions that were significantly different between the groups, which were enriched in 4 metabolic pathways. Wilcoxon signed-rank test of COBRA-predicted metabolic fluxes revealed significant differences in the fluxes of 21 metabolites. Although the abundance of six species did not differ significantly between the two groups, their contributions to metabolic fluxes were significantly different. Mediation analysis indicated that gut microbiota influence the progression of depression by modulating various metabolites.

**Conclusions:** This study combined species abundance with COBRA metabolic flux predictions and mediation analysis, identified several differential species, as well as multiple differential metabolic fluxes, such as Cu^2+^ and L-Dopa, which were attributed to species like *Dorea*. Species such as *Acholeplasma* and *Acidovorax* were found to influence the progression of depression by affecting various metabolites, including Cu^2+^. These findings contribute to a deeper understanding of the relationship between gut microbiota and depression, offering new directions for the diagnosis and treatment of depression.

## INTRODUCTION

Depression is one of the most prevalent mental disorders, characterized primarily by a prolonged low mood, high prevalence, recurrence rates, and suicidal tendencies^1^. Despite its profound social and individual impacts, the pathogenesis of depression remains poorly understood and presents substantial challenges for diagnosis and treatment.

The human gut contains a large number of microorganisms, including bacteria, viruses, archaea, and fungi. The proteins and metabolites they encode can significantly influence human health^2^,^2^. Research indicates that the gut microbiota has the potential to interact with the central nervous system through various pathways, potentially influencing the onset and progression of depression^4–6^. During the past decade, increasing evidence has suggested that the brain-gut-microbiota axis plays a crucial role in depression. Causal studies suggest that alterations in the gut microbiota contribute to the development of depression and raise its risk^7^.

An increasing number of studies suggest a potential link between gut dysbiosis and depression. Although results vary, a common finding is an increase in pro-inflammatory bacteria and a decrease in anti-inflammatory bacteria^6^,^8^. In depression, the phyla Firmicutes, Actinobacteria, and Bacteroidetes are the most affected^9^,^10^, particularly with an increase in the Bacteroidetes/Firmicutes ratio. Previous studies have shown that several species of the Bacteroides genus (e.g., *B. thetaiotaomicron, B. fragilis*, and *B. uniformis*) significantly increase the risk of depression^11^,^12^.

Changes in the composition of gut microbiota in patients with depression can lead to alterations in the microbial metabolome^13^. Transplantation of fecal microbiota from individuals with depression into germ-free mice can induce depressive-like behaviors in the recipients^9^. These mice predominantly exhibit disruptions in microbial genes and host metabolites, particularly those involved in carbohydrate and amino acid metabolism^9^. Van et al. have shown that retinol (vitamin A), 4-hydroxycoumarin, 2-aminooctanoate, 10-undecenoate, 1-palmitoyl-2-palmitoleoyl-GPC, and 1-linoleoyl-GPA are associated with depression^13^, and hippurate is considered a biomarker for depression^14^. Skonieczna et al. found that short-chain fatty acids (SCFAs), common microbial metabolites in the gut, are closely associated with depression, exhibiting significantly lower concentrations in the feces of depressed patients compared to healthy individuals^15^.

Due to the sparsity of microbiome data and significant inter-individual variability^2^,16–18, directly identifying microbial features associated with depression is challenging. Microbial metabolites, as the final products of microbial genomes, directly influence the gut environment, making metabolomic data a more accurate reflection of the gut microbiota-depression relationship. Therefore, this study uses species abundance data and applies constraint-based reconstruction and analysis (COBRA) methods^19^,20 to predict gut microbial metabolic fluxes in both depression patients and healthy controls, analyzing microbiota and metabolic dysbiosis in depression. By examining how different species contribute to dysregulated metabolic flux in depression patients and conducting mediation analysis, we aim to gain a deeper understanding of the connection between gut microbiota and depression.

## Materials and Methods

### Data Collection and Depressive-Healthy Pair Matching

The samples used in this study were sourced from the GMrepo v2 database (https://gmrepo.humangut.info/home)^21^. A total of 354 depression samples and 5575 healthy samples were collected (Figure 2).

**Figure 1.**
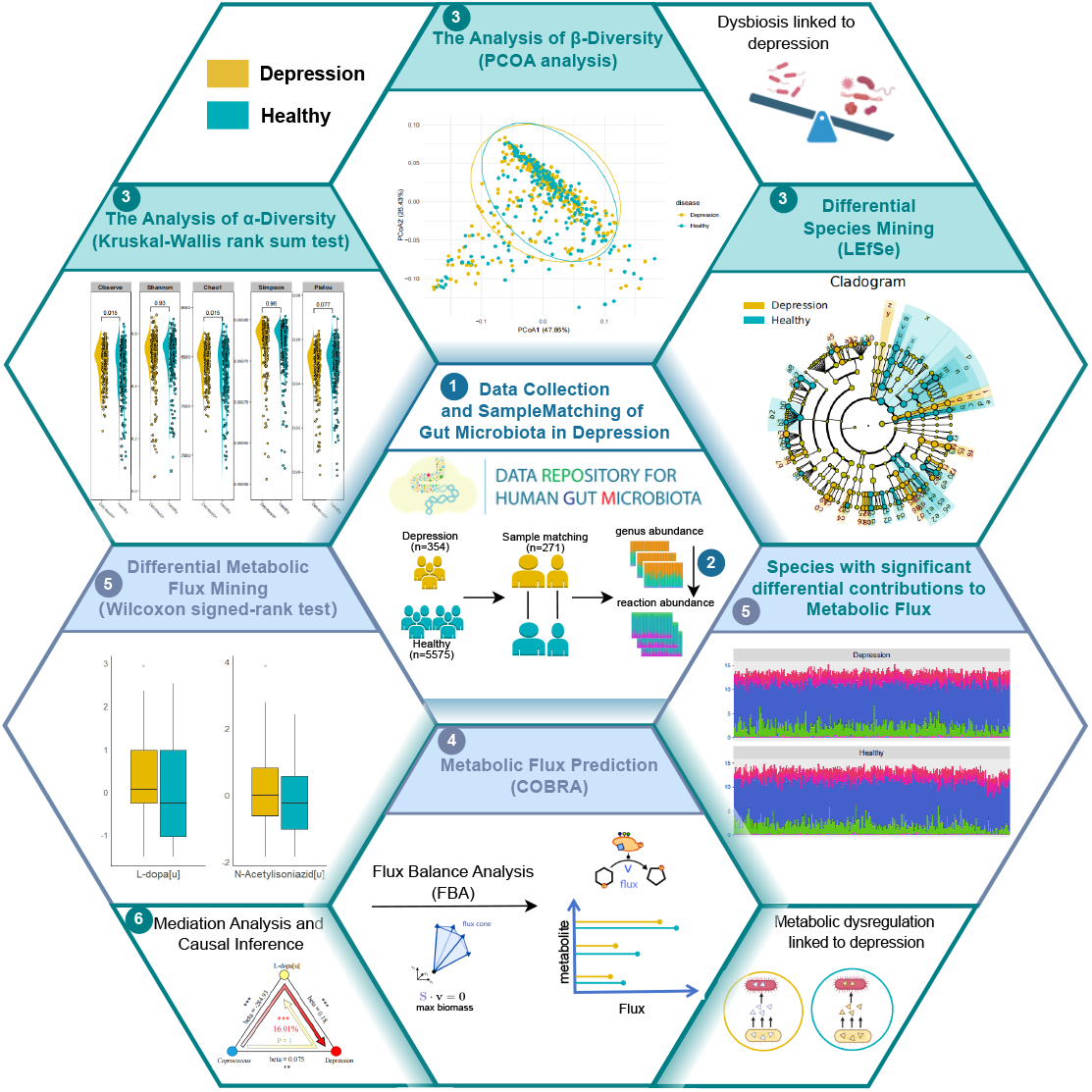
The overview of this study. Step 1: Collect samples from the GMrepo v2 database and perform matching. Step 2: Convert species abundance data into reaction abundance. Step 3: Conduct diversity analysis on species abundance and reaction abundance, and identify differential species using LEfSe. Step 4: Perform flux balance analysis (FBA) for metabolic flux prediction. Step 5: Conduct differential testing of metabolic fluxes and analyze the contributions of different species to metabolite flux. Step 6: Mediation analysis of species, metabolites and depression.

**Figure 2.**
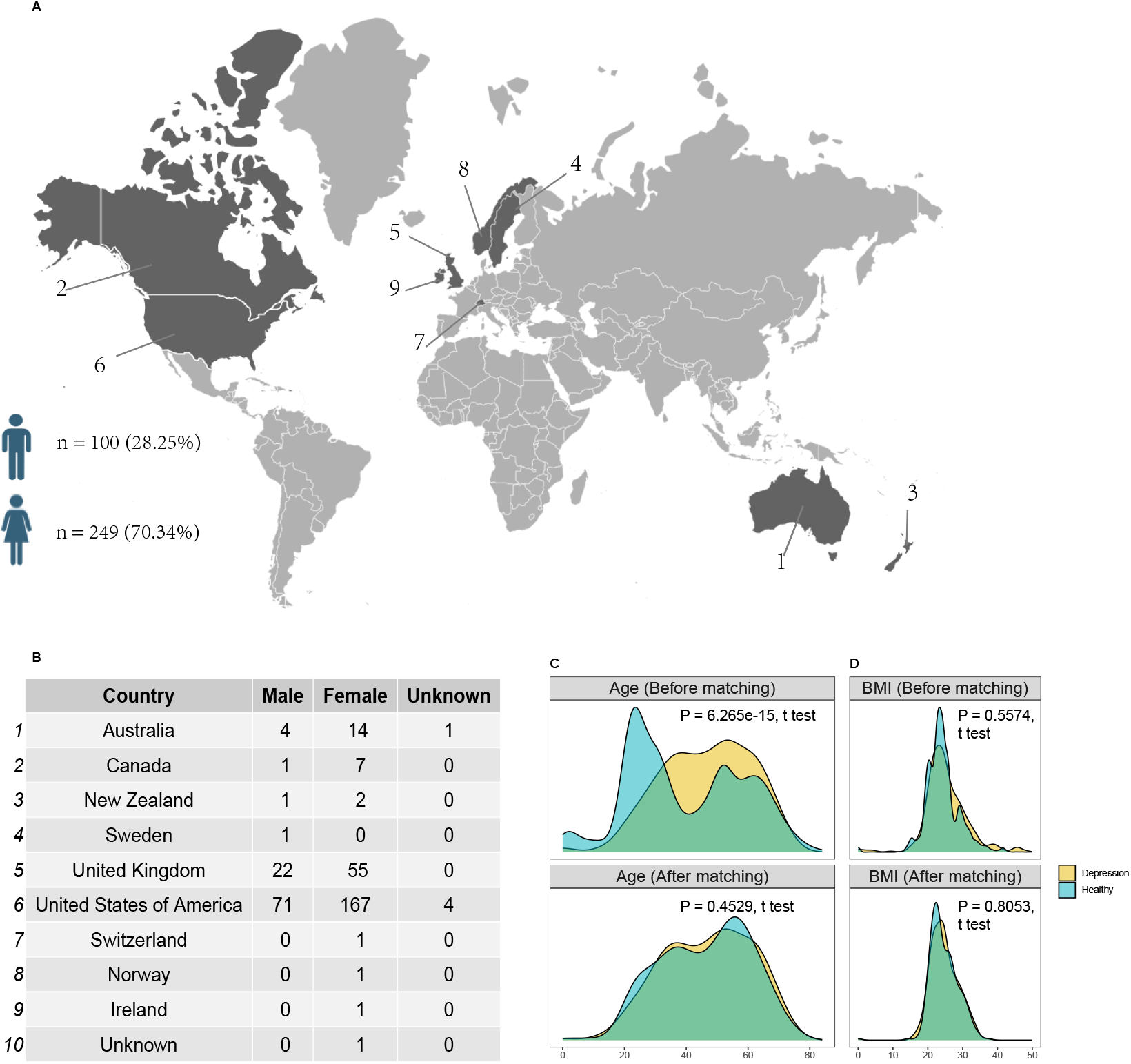
Nationality, gender, age and BMI distributions before and after matching for the samples. A,B. The countries of samples; C,D. Age distribution before and after matching between depression patients and healthy individuals (P = 6.265e-15 and 0.5574, t test). E, F. BMI distribution before and after matching between depression patients and healthy individuals (P = 0.5574 and 0.8063, t test).

To minimize the effects of potential confounding factors, such as age and BMI, that could lead to misleading results, we performed depressive-healthy pair matching. This process ensures that observed effects are accurately attributed to the independent variables under investigation, rather than confounding variables^22^. We developed an algorithm to match healthy samples with depression samples based on different variable types:

- **Categorical Variables (e.g**., **nationality, gender)**: We match healthy samples exactly with depression samples on these variables.
- **Numerical Variables (e.g**., **age, BMI)**: To reduce the influence of extreme values, we calculated the 25th and 75th percentiles (the interquartile range) and computed the standard deviation within this range. The healthy samples were then selected if their values for these variables were within one standard deviation of the depression sample.

Next, we performed sample matching, resulting in several possible outcomes: (1) no match, (2) one-to-one match, (3) one-to-many match, or (4) many-to-one match. To maximize the number of matched pairs, we converted the results into a bipartite graph and applied the Hungarian algorithm^23^,24 for maximum bipartite matching, ensuring the largest possible number of valid matches (Algorithm 1).

### Species Abundance Processing

Microbial relative abundance data are compositional, meaning the proportions of all species in a sample sum to 1. This inherent dependency between species makes it unsuitable for statistical methods that assume variable independence. To address this, we applied the centered log-ratio transformation (CLR)^25^, which eliminates interdependence and makes the data compatible with conventional statistical analyses. The CLR transformation also enhances the detection of patterns and relationships within the data, enabling more accurate interpretation and comparison.

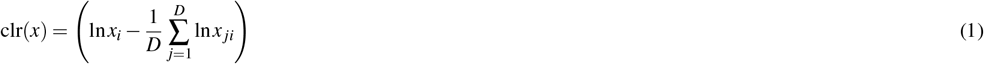

Where *x*_*i*_ is the relative abundance of the *i*-th species, and *D* denotes the total number of species. The CLR transformation calculates the natural logarithm of the abundance of each species relative to the geometric mean of all species’ abundances, effectively normalizing the data to address the compositional nature of microbiome datasets.

#### Algorithm 1 Depressive-Healthy Pair Matching Algorithm

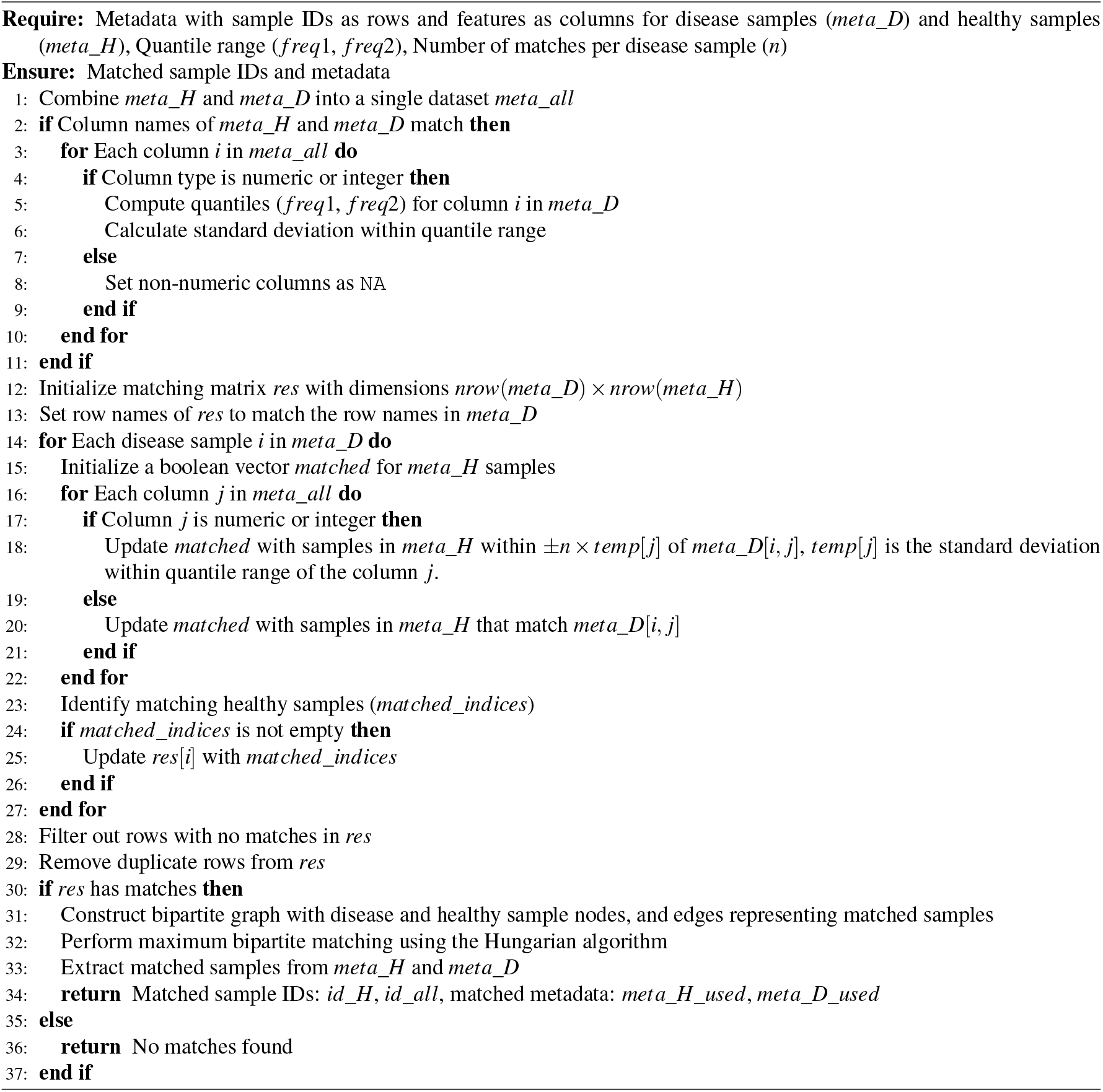

### Diversity and differential analysis

To evaluate gut microbial diversity differences between depression patients and healthy individuals, we calculated several alpha diversity indices, including the Pielou, Shannon, and Simpson indices. The nonparametric Kruskal-Wallis rank sum test was used to assess statistical differences in diversity between the two groups. Beta diversity analysis was performed to examine differences in species composition. Principal coordinate analysis (PCoA) was used to visualize differences between microbial communities, while the ANOSIM test was applied to statistically compare group similarities. Additionally, LEfSe (Linear Discriminant Analysis Effect Size) was employed to identify species significantly associated with depression or health status.

### Metabolic flux prediction and reaction abundance calculation

#### Constructing a generic species model at the genus level

The AGORA2 database^26^ contains metabolic models for 7,302 different strains, detailing all known metabolic reactions in each strain, including their stoichiometry and directionality^27^,^28^. These models are derived from genome data, annotations, and strain-specific biochemical and physiological literature^28^. When constructing a genus-level model, all metabolic reactions of strains within the genus were consolidated into a single model after removing duplicates. The biomass reaction for the genus was represented by the average biomass reaction of all strains. Metabolic models represent strain growth states using a biomass reaction, which includes processes associated with macromolecules such as nucleic acids and proteins. These models can be visualized as networks, where edges represent connections between metabolites enabled by specific reactions (Figure 3).

**Figure 3.**
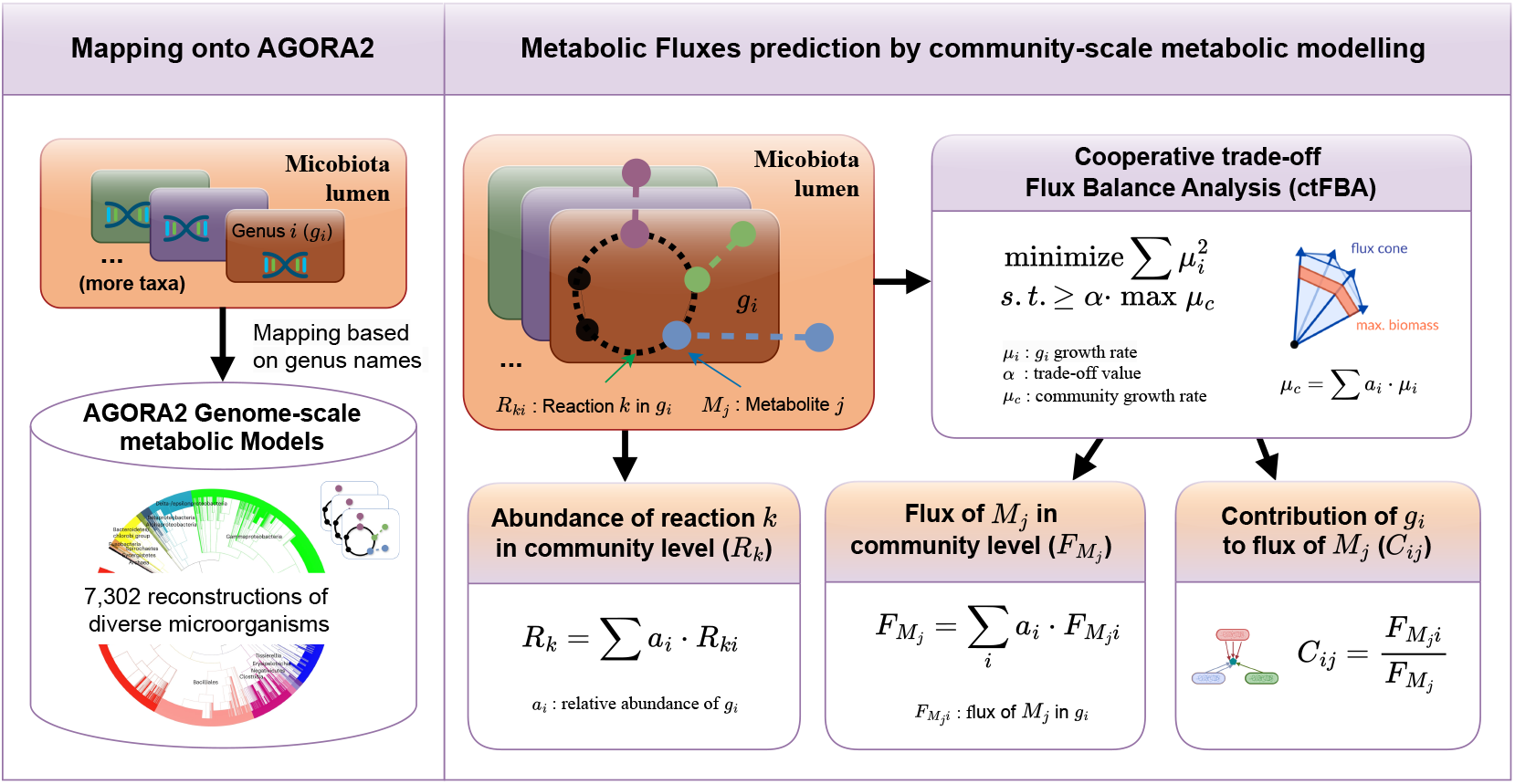
Workflow of metabolic flux analysis using AGORA2. The process includes flux balance analysis (FBA) to generate a flux matrix, reconstruction of the metabolic network, calculation of reaction abundance, and determination of species contributions to metabolic flux. In this workflow, Here, *G* represents the genus, *m* denotes the number of genera, *R* is an *n ×* 1 vector indicating the presence of each reaction in a genus, where *n* is the total number of reactions. *R*_*c*_ represents the reaction abundance of the community, *a* is the abundance of the genus, *M*_*i, j*_ represents the *j*-th metabolite in the *i*-th species, and *k* denotes the total number of metabolites.

#### Constructing individualized gut microbiota metabolic models

Individualized gut microbiome metabolic models were constructed using the mgPipe workflow from the COBRA Tool-box^29^.

#### FBA (Flux Balance Analysis) for predicting metabolic reaction rates

Flux Balance Analysis (FBA) is a widely used method for analyzing metabolic networks^20^. It predicts the steady-state flux distribution of reactions within a metabolic network under given conditions by solving a linear programming problem (Figure 3).

#### Calculating the Reaction Abundance

To calculate the reaction abundance in each community model: 1. Combine all reactions across the genus in the community into a single 1\**n* vector *R*, where *n* is the total number of reactions, *m* is the total number of genus. 2. Compute the communitys reaction abundance *R*_*c*_ using the formula:

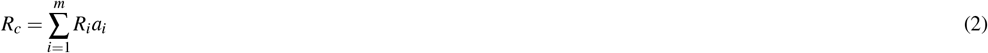

Where: - *R*_*i*_: A binary vector for genus *i*, where each column is 1 if the reaction exists in the genus and 0 otherwise. - *a*_*i*_: The relative abundance of genus *i*.

Thus, *R*_*c*_ is the weighted sum of the reaction presence vectors *R*_*i*_, with weights corresponding to the relative abundances of each genus. This approach quantifies the functional contribution of each genus to the overall metabolic activity of the community.

### Statistical Analysis

#### Differential and Enrichment Analysis

To compare differences between two paired groups, the Wilcoxon signed-rank test was applied^30^. For enrichment analysis, the hypergeometric test was employed^31^ was employed. To explore how gut microbiota influence depression progression through metabolites, we performed mediation analysis using the R package mediation. All statistical analyses were carried out using R version 4.2.0.

## RESULTS

### Balanced Depressive-Healthy Cohort through Sample Matching

To reduce the influence of confounding factors, sample matching was performed, resulting in 271 depression samples alongside their matched healthy controls. In the original dataset, significant differences existed in the distribution of age and BMI between the depression and healthy groups; however, following matching, these differences were substantially diminished(Figure 2). The mapping of genus-level species names with the AGORA2 database shows that all genus-level species in the depression samples can be matched to AGORA2, and the genus-level species coverage in the healthy control samples also reaches over 60% (Supplementary Figure 2). This indicates that the metabolomic model dataset constructed in this study has good species coverage, allowing for a comprehensive and accurate representation of the overall metabolic characteristics of the microbiome, thereby providing a reliable foundation for subsequent in-depth metabolic network analysis.

### Microbiota Dysbiosis in Depression

The α-diversity indices for depression and healthy samples was calculated (Figure 4A). The Observed (P < 2.2e-16, Wilcoxon rank-sum test), Shannon (P = 0.045, Wilcoxon rank-sum test) and Chao1 (P < 2.2e-16, Wilcoxon rank-sum test) indices showed significant differences between the two groups, indicating that the richness and evenness of the gut microbiota were slightly reduced in depression patients. β-diversity analysis revealed that there were small but significant differences in gut microbiota composition (Figure 4B) between depression patients and healthy individuals (R = 0.008, P = 0.004, ANOSIM test). To further explore these differences, LEfSe analysis was performed. The LEfSe analysis identified 18 species (LDA score > 2 and P < 0.005) significantly enriched in healthy samples and 5 species enriched in depression samples (Figure 4C). At the genus level, 16 genera exhibited differences, while 5 species differed at the family level, and 1 species each at the class and order levels. Figure 4D shows the taxonomic classification diagram of the 23 species.

**Figure 4.**
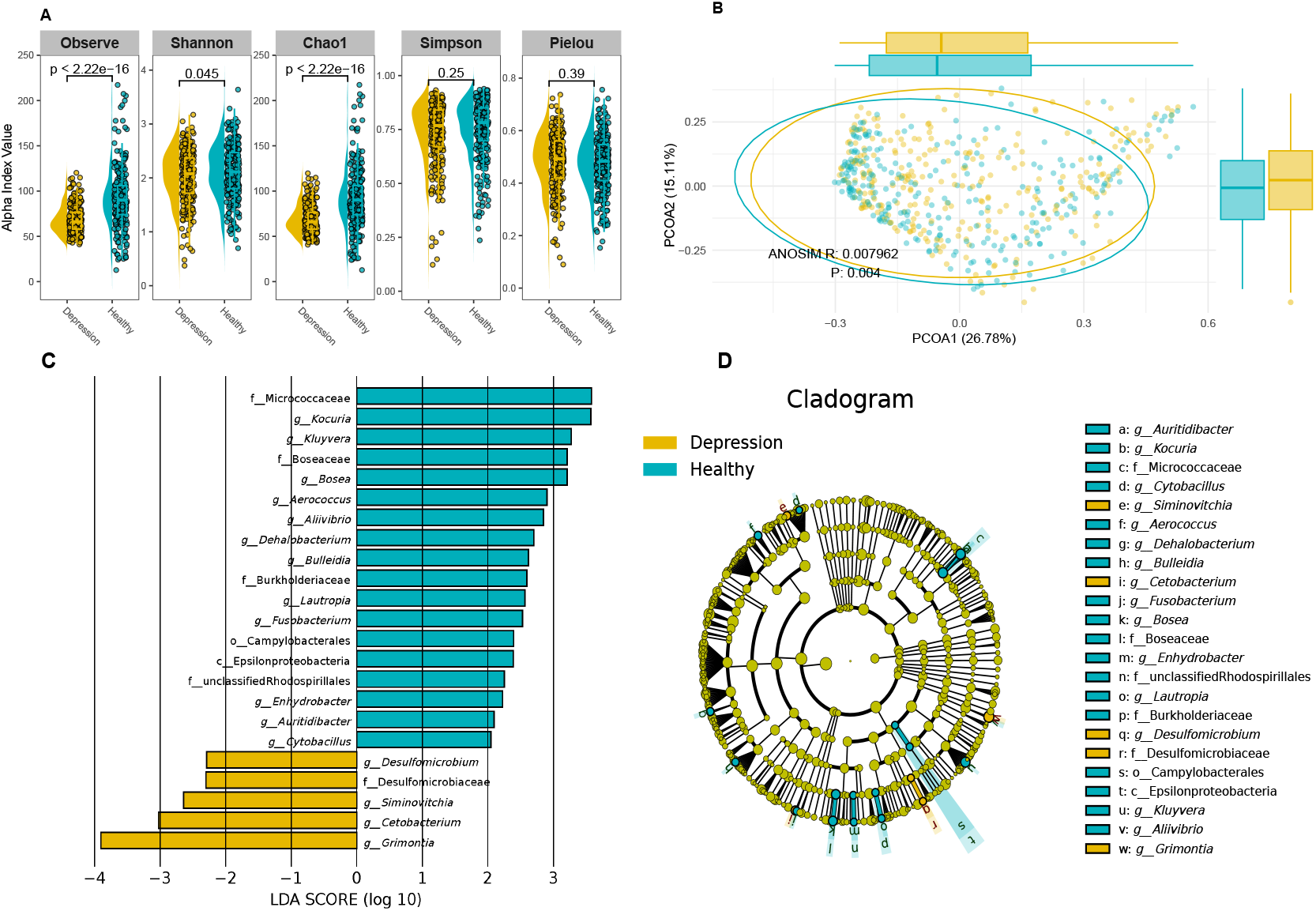
Species diversity analysis. A. The difference of alpha diversity indices between depression patients and healthy individuals.(Observed: P < 2.2e-16, Shannon: P = 0.045; Chao1: P < 2.2e-16) B. The differences in community composition between the depression patients and healthy individuals. (ANOSIM R = 0.008, P = 0.004). C. The 23 differential species between depression and healthy samples(LDA score > 2 and P < 0.005), with 5 species enriched in depression samples and 18 species enriched in healthy samples. D. Taxonomic classification diagram of significantly different species.

### Metabolic Reaction Dysregulations in Depression

By integrating species abundance, reaction abundance, and pathway abundance into a 0-1 matrix (Supplementary Fig. 1), it is evident that species sparsity is notably high in both depression and healthy samples, indicating that species-level markers may lack generalizability for identifying universal markers of depression. In contrast, the composition of metabolic reactions and subsystems showed greater consistency across samples, suggesting that these levels may hold more robust and universal markers for depression, warranting deeper investigation.

Alpha diversity analysis of reaction abundance showed significant differences between groups for the Observed (P = 0.015, Wilcoxon rank-sum test) and Chao1(P = 0.015, Wilcoxon rank-sum test) indices (Figure 5A). However, beta diversity analysis found no significant differences in reaction abundance between the groups (R:-0.0003, P:0.493, ANOSIM test, Figure 5B).

**Figure 5.**
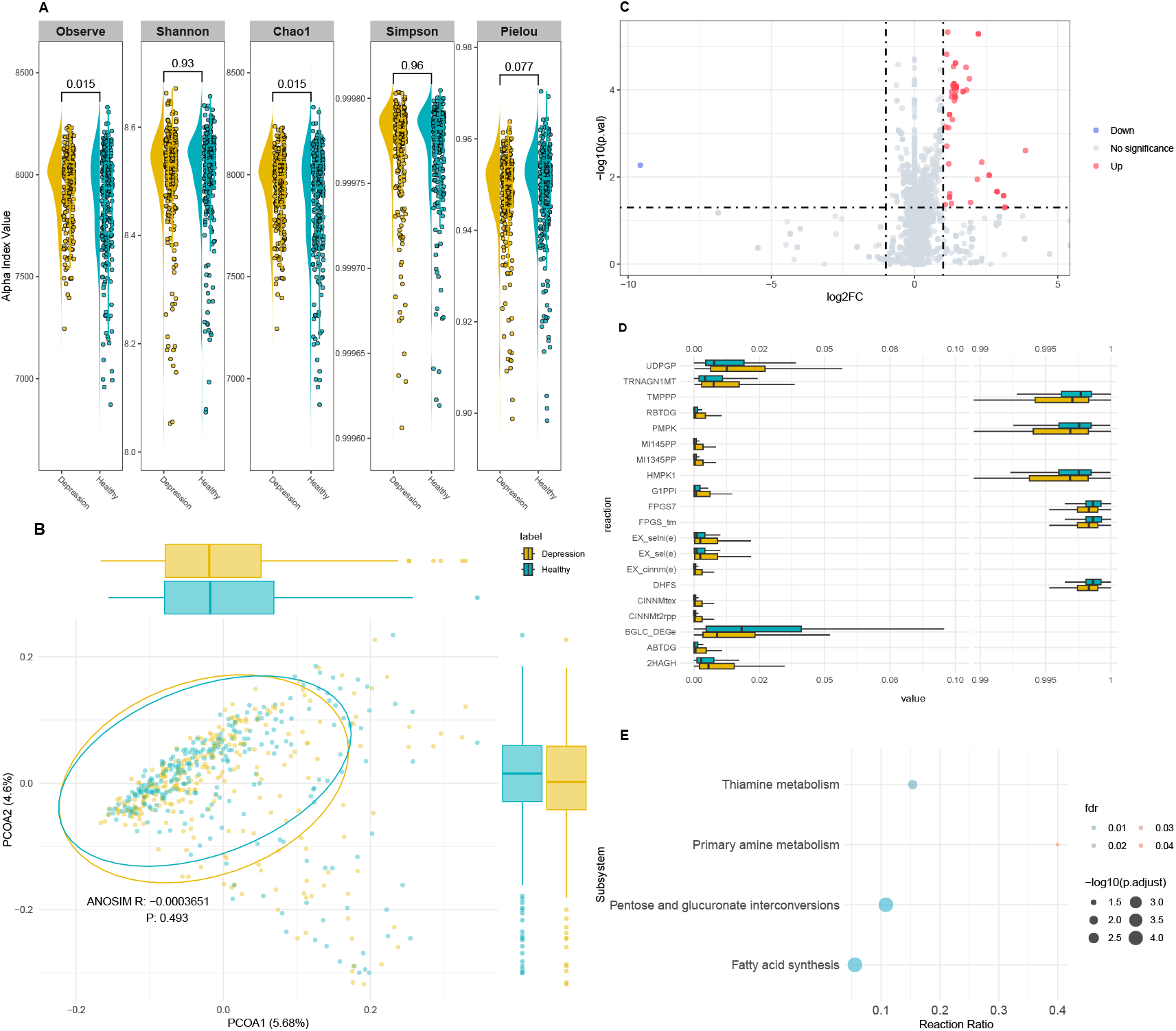
Metabolic reaction diversity analysis. A. The difference of alpha indices in reaction abundance between depression patients and healthy individuals. (Observed: P = 0.015; Chao1: P = 0.015). B. There is no differences in reaction composition between the depression patients and healthy individuals (ANOSIM R: -0.0003, P: 0.493). C. 89 differential reactions were identified (FC > 2 and P < 0.05). D. Top 20 reactions with the most significant differences between the two groups (P < 0.05, Wilcoxon signed-rank test). E. Enrichment of 89 significantly different reactions into 4 subsystems. The size of the points represents the number of different reactions within each subsystem, and the color represents the FDR value.

To identify specific reactions with differences, the Wilcoxon signed-rank test revealed 89 significantly altered reactions (FDR < 0.05, Figure 5C). The top 20 reactions with the smallest FDR values are shown as boxplots in Figure 5D. Enrichment analysis of these 89 reactions identified four subsystems with significant enrichment: Fatty acid synthesis, Pentose and glucuronate interconversions, Thiamine metabolism, and Primary amine metabolism (FDR < 0.05, Figure 5E).

### Metabolic Flux and Species Contribution Differences

To obtain the metabolic fluxes, genus-level metabolic models were solved using flux balance analysis (FBA) with the objective of maximizing the community biomass reaction. A Wilcoxon signed-rank test identified 21 metabolites with significantly different fluxes between depression patients and healthy individuals (FDR < 0.05, Figure 6A). Among the 21 differential metabolites identified, 13 were recorded in The Human Metabolome Database (HMDB). Among these 13 metabolites, 1 belongs to Homogeneous transition metal compounds, 2 to Non-metal oxoanionic compounds, 1 to Carboxylic acids and derivatives, 1 to Fatty Acyls, 1 to Imidazopyrimidines, 3 to Organonitrogen compounds, 1 to Organooxygen compounds, 2 to Purine nucleotides, and 1 to Steroids and steroid derivatives.

**Figure 6.**
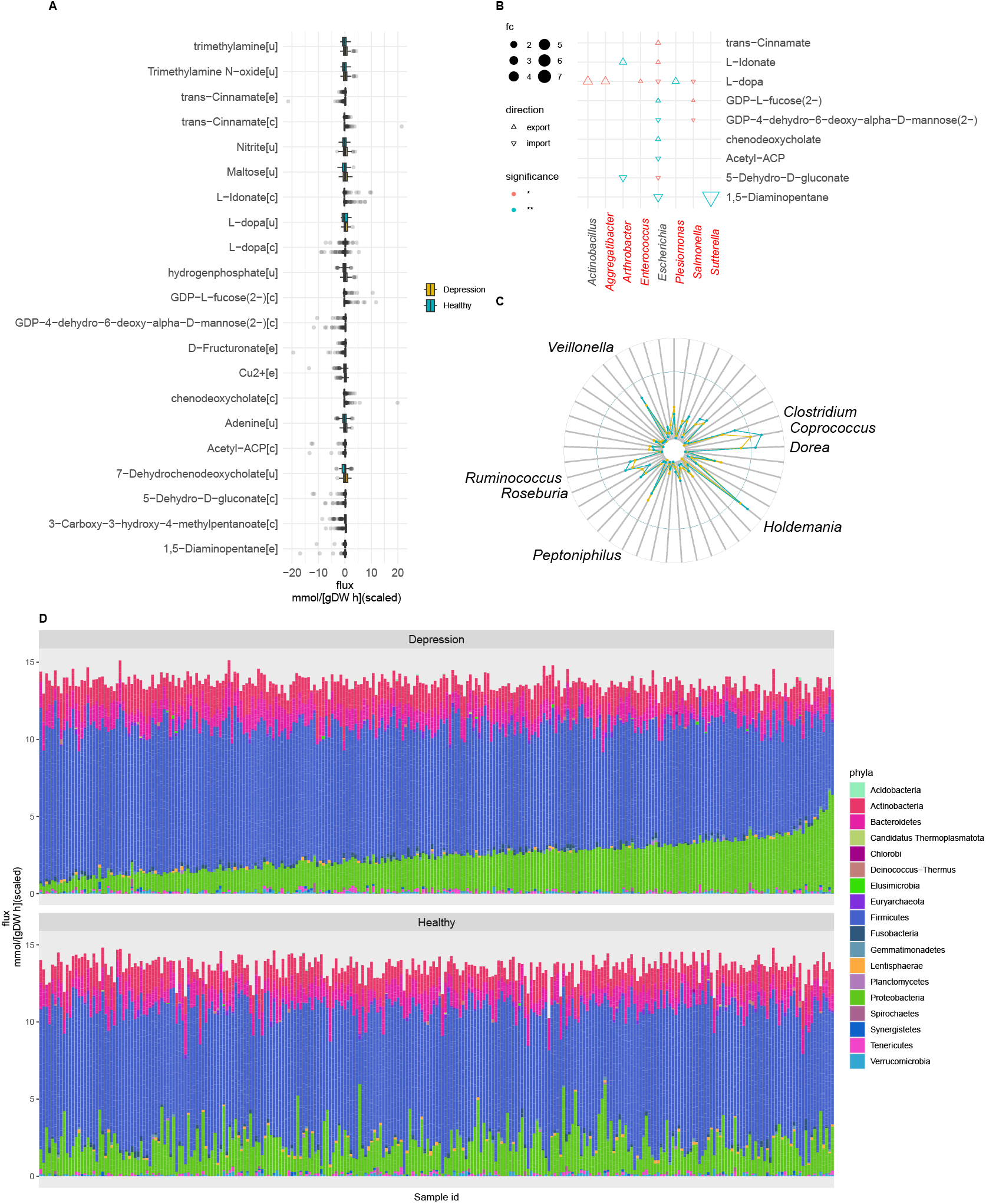
Significant differential metabolic Fluxes. A. Significant differences in metabolic flux between depression patients and healthy individuals (P < 0.05, Wilcoxon signed-rank test). [u]:thylakoid, [e]: extracellular space, [c]: cytosol. B. Species with significant differential contribution to metabolic flux, with dot size representing fold change (FC) and color indicating P-values. Species highlighted in red show no abundance difference between the two groups, but exhibit significant differences in their contribution to metabolic flux. C,D. Cu^2+^ consumption flux differences in genus(C) and phyla(D) between the two groups. The top 8 genera with the highest contributions are displayed in text in C.

Further analysis of species contributions to the metabolites revealed that 6 species, despite no significant differences in their abundance between the groups, exhibited significant differences in their contributions to metabolite fluxes (Figure 6B). Notable metabolites such as L-Dopa and *Cu*^2+^ were highlighted. The results revealed that 6 genera showed significant differences in their contributions to L-Dopa metabolism between depression patients and healthy individuals (Figure 6B). Among these, 4 genera (highlighted in red) exhibited no significant differences in abundance between the two groups. Analysis of *Cu*^2+^ metabolism revealed that species such as *Dorea* and *Coprococcus* have a significantly higher rate of *Cu*^2+^ consumption in depression patients compared to healthy individuals (Figure 6C). Figure 6D illustrates the phylum-level contributions to *Cu*^2+^ metabolism, highlighting distinct differences between the two groups.

### Microbiota-Depression Causal Inference via Mediation Analysis

Mediation analysis was conducted to assess how much metabolites mediate the influence of gut microbes on depression. This analysis identified 9 significant mediation linkages (Figure 7A, all *P*_*mediation*_ < 0.001, *P*_*inversemediation*_ = 1). These linkages involved 4 genera (*Coprococcus, Escherichia, Papillibacter*, and *Ruminococcus*) and 8 metabolites, including Cu^2+^ and L-Dopa.

**Figure 7.**
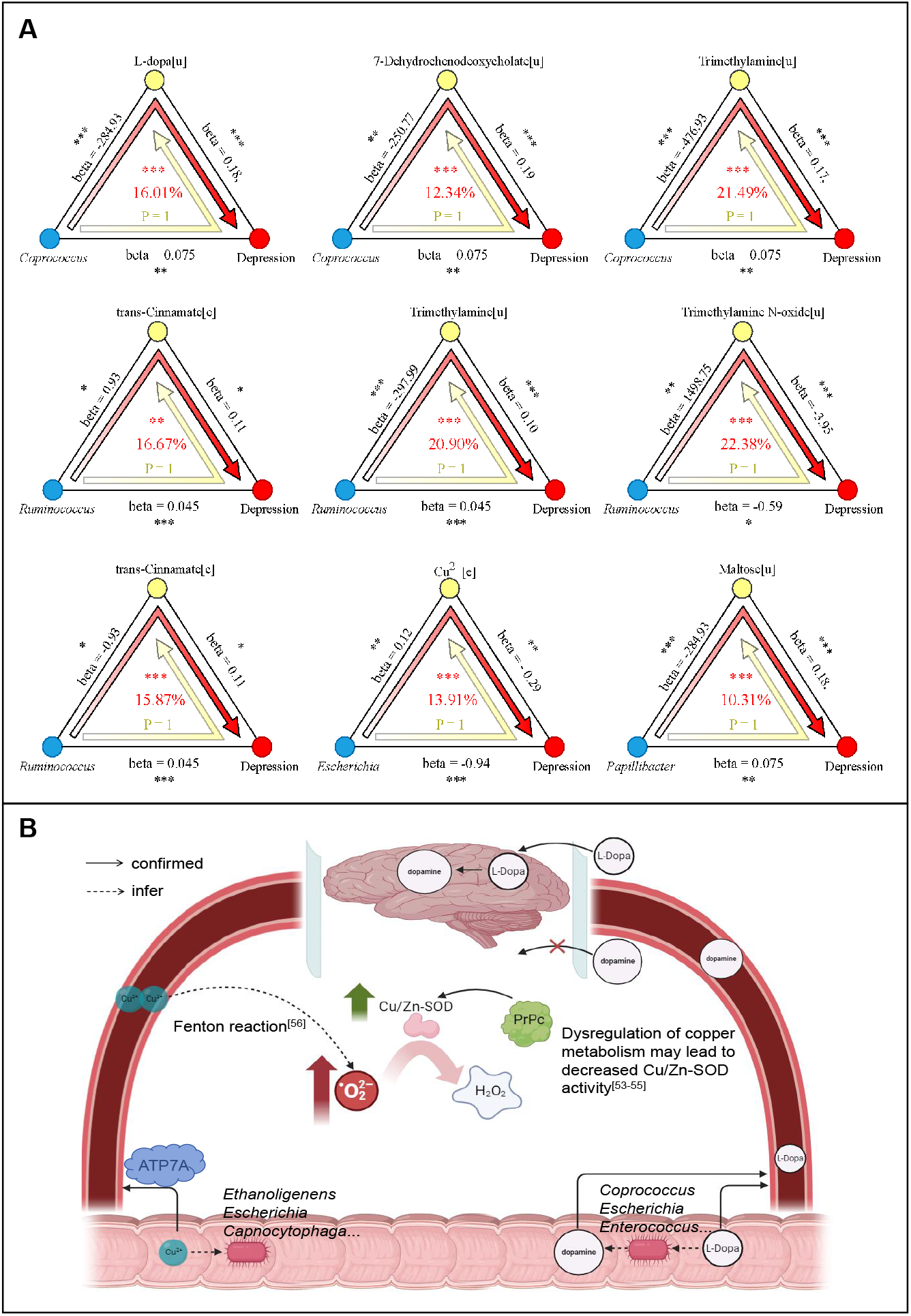
Causal mediation analysis to explore the mediating effect of microbial metabolites on depression. A. Nine causal pathways linking species, metabolites and depression. B. Inferred causal mechanisms of gut microbiota in depression, solid arrows represent known paths, while dashed arrows represent inferred paths. Left: Reduced Cu^2+^ consumption by gut microbes increases Cu^2+^ concentration in the bloodstream, promoting oxidative stress and raising the risk of depression. Right: Gut microbes metabolize L-Dopa into dopamine, reducing the availability of L-Dopa, which can cross the blood-brain barrier, for brain uptake, thereby increasing the risk of depression.

The findings suggest distinct mechanisms underlying these relationships: Gut microbial activity reduces Cu^2+^ consumption, leading to elevated systemic Cu^2+^ levels. This increase exacerbates oxidative stress in the host, potentially accelerating the progression of depression. As a precursor to dopamine, L-Dopa can cross the blood-brain barrier and is converted to dopamine in the brain. However, in individuals with depression, gut microbes metabolize L-Dopa into dopamine within the gut, reducing its availability for brain uptake and disrupting the dopamine pathway (Figure 7B).

## DISCUSSION

This study explores the complex relationship between gut microbiota, metabolic reactions, and depression. Our findings suggest that while species diversity (α-diversity) and overall microbial composition (β-diversity) shows limited differences between depression and healthy groups, metabolic reactions and metabolites may offer a more comprehensive understanding of depression-related biomarkers.

Several genera, including *Auritidibacter* and *Kocuria*, were newly associated with depression in this study (Table1). *Auritidibacter*, known for its association with ear infections^32^,^33^, and *Kocuria*, a genus with limited prior research, represent intriguing candidates for further investigation due to their novel link to depression. Conversely, taxa such as *Micrococcaceae* and *Burkholderiaceae*, previously reported as enriched in depression^34^,^35^, were found to be enriched in healthy individuals in this study, raising questions about their relevance as depression-associated markers.

**Table 1.**
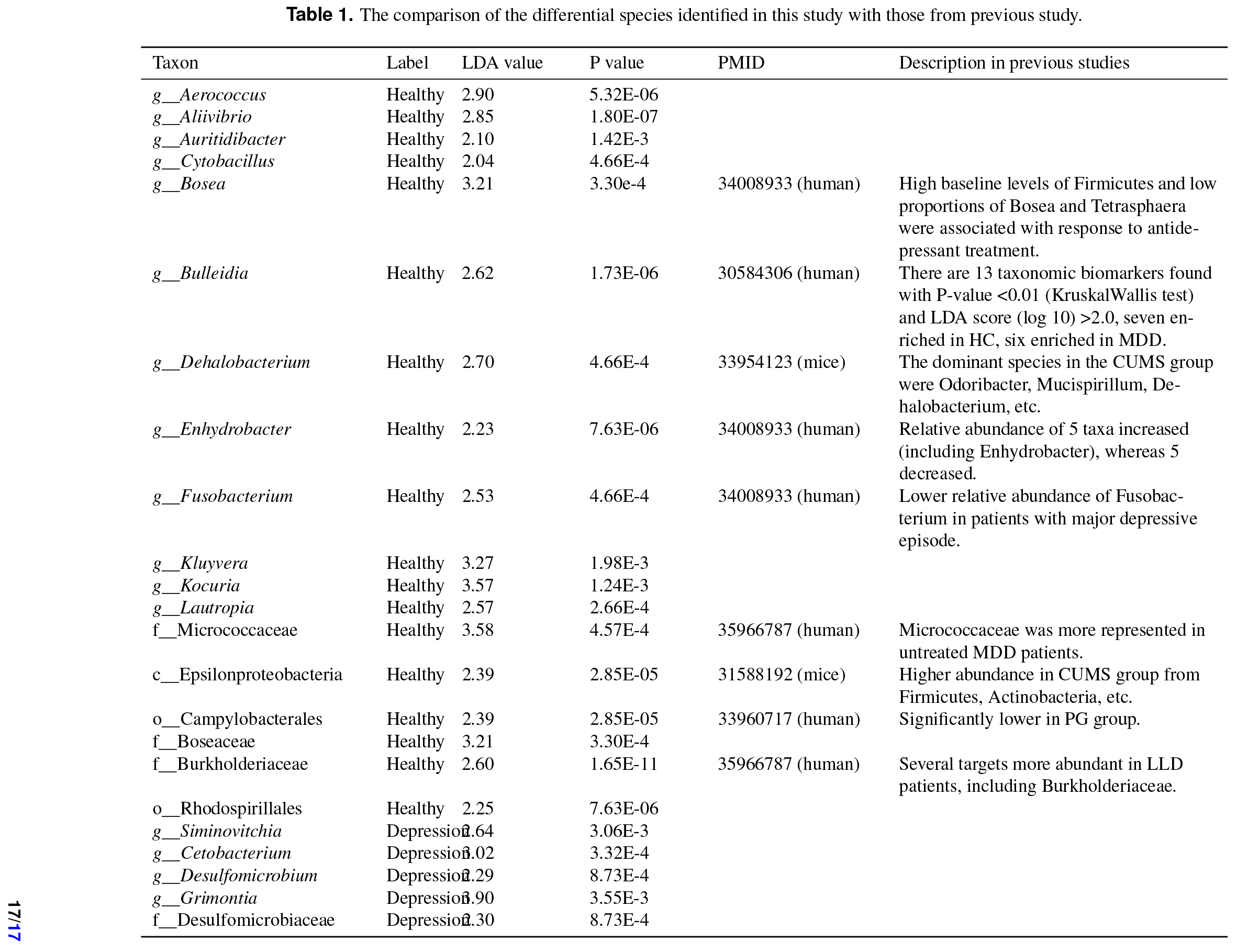
The comparison of the differential species identified in this study with those from previous study.

Unlike the sparse species diversity, metabolic reactions and metabolites were consistently present across samples, providing a more stable foundation for identifying depression-related biomarkers. Metabolic flux analysis revealed 89 significantly altered reactions enriched in four metabolic pathways: fatty acid synthesis, pentose and glucuronate interconversions, thiamine metabolism, and primary amine metabolism. These findings point to metabolic processes as critical factors in depression pathophysiology.

One particularly intriguing finding relates to copper (Cu) metabolism, where Cu plays a crucial role in oxidative stress regulation as a cofactor for antioxidant enzymes such as Cu/Zn superoxide dismutase (Cu/Zn-SOD). This enzyme converts harmful superoxide anions into hydrogen peroxide, mitigating free radical-induced cellular damage. In depression, oxidative stress levels are elevated, and antioxidant enzyme activity is reduced, with the severity of depression positively correlated with oxidative stress markers^36^. Dysregulation of copper metabolism may reduce Cu/Zn-SOD activity (Figure 7B), exacerbating oxidative damage and contributing to the pathophysiology of depression^37–39^. The brain is particularly vulnerable to oxidative stress due to its high levels of unsaturated fatty acids, elevated oxygen consumption, and limited antioxidant defenses^40^, which can lead to neuroinflammation and cognitive deficits. Studies have found significantly higher blood Cu levels in patients with depression compared to healthy individuals^41^. This excess Cu, possibly stemming from reduced Cu consumption by gut microbiota, disrupts copper homeostasis and catalyzes the Fenton reaction, producing excessive reactive oxygen species (ROS) and amplifying oxidative stress^42^. Elevated Cu levels have also been linked to memory impairment by disrupting the expression of hippocampal proteins like GluN2B and PSD95, critical for synaptic function. Furthermore, dietary intake of micronutrients such as zinc, iron, copper, and selenium shows a negative correlation with depression, indicating a protective role for balanced micronutrient intake. The copper-binding PrPc protein enhances SOD activity and reduces oxidative stress, with PrPc-deficient mice displaying depression-like behaviors, further highlighting the link between copper regulation and depression (Figure 7B).

L-Dopa is the natural form of dihydroxyphenylalanine and the direct precursor of dopamine. Unlike dopamine, L-Dopa can crosses the blood-brain barrier and is metabolized into dopamine by aromatic L-amino acid decarboxylase. This study identified 6 genera with significant differences in their contributions to L-Dopa between depression patients and healthy individuals. Among the nine metabolites shown in Figure 6B, *Enterococcus* exhibited a significant difference only in its contribution to L-Dopa. The genus *Enterococcus* plays a key role in L-Dopa metabolism in the gut^43^, converting L-Dopa into dopamine via tyrosine decarboxylase. This process reduces the amount of L-Dopa available for brain uptake, potentially impacting depression pathology.^44–46^. A previous study indicated that the level of L-Dopa in the intestines of chickens treated with recombinant attenuated Salmonella vaccine was increased^47^; however, no association between L-Dopa metabolism and the remaining species has been reported. L-Dopa is metabolized into dopamine by gut microbes, thereby reducing the brain’s utilization of L-Dopa(Figure 7B)

Moreover, our findings emphasize the importance of gut microbiota in modulating metabolic processes. Certain microbial species, despite showing no significant differences in abundance, demonstrated markedly distinct contributions to specific metabolic pathways, such as those involving L-Dopa and *Cu*^2+^. These species, though low in abundance, exert substantial effects on host metabolism, suggesting that even minor microbial shifts can have profound effects on host physiology.

In summary, our results underscore the value of examining both microbial composition and metabolic reactions in understanding depression. While microbial species alone may not serve as reliable markers, the broader metabolic network, influenced by microbial activity, offers a more stable and meaningful perspective on depression’s underlying mechanisms. Future research should focus on these metabolic pathways and their modulation by the microbiota to identify novel diagnostic and therapeutic targets for depression.

## CONCLUSION

In this study, constraint-based metabolic modeling was utilized to explore the relationship between the gut microbiome and depression. Significant differences in microbial composition and diversity were observed between depression patients and healthy individuals, with specific bacterial genera such as Auritidibacter and Kocuria enriched in the depression group. Metabolic pathway analysis identified notable alterations in fatty acid synthesis, pentose and glucuronate interconversions, thiamine metabolism, and primary amine metabolism. Furthermore, depression patients exhibited a reduced capacity for L-Dopa metabolism, potentially affecting dopamine levels, and lower Cu^2+^ consumption by gut bacteria, suggesting increased oxidative stress. These findings highlight the potential role of gut microbiota in depression pathophysiology and offer novel insights for developing microbiota-targeted therapeutic strategies.

## KEY POINTS

- Gut Microbiome Diversity and Depression: The study reveals that individuals with depression exhibit significantly lower gut microbial diversity compared to healthy controls. This reduction is particularly evident in species richness and evenness, with certain species like those in the Micrococcaceae family enriched in depressed individuals, while Firmicutes are more abundant in healthy controls.
- Metabolic Pathway Alterations: Significant differences were identified in metabolic pathways between the depression and control groups. These include fatty acid synthesis, pentose and glucuronate interconversions, thiamine metabolism, and primary amine metabolism. The findings suggest that these altered metabolic functions in the gut microbiota may contribute to the onset and progression of depression.
- Potential Biomarkers and Therapeutic Targets: The study identifies 21 metabolites with significant differences between the two groups, with L-Dopa metabolism and Cu^2+^ metabolism standing out. These metabolic changes point to potential biomarkers for diagnosing depression and offer novel therapeutic targets, such as modulating gut microbiota to impact mental health.

## Supporting information

https://github.com/zyc134/Depression

## Data availability

All data is available in the GMrepo V2 database (https://gmrepo.humangut.info/home). The data and code used in this study can be accessed in the GitHub repository at https://github.com/zyc134/Depression

